# Incidence and Risk Factors of Combined-Antiretroviral Therapy-Induced Hepatotoxicity among HIV Patients at the Bali District Hospital, Cameroon

**DOI:** 10.1101/2020.11.16.384339

**Authors:** Yayah Emerencia Ngah, Frederick Nchang Cho, Bisong Shauna Etagha, Neh Gladys Fusi, Neba Francisca, Mondinde George Ikomey, Njimona Ibrahim

**Affiliations:** Dapartment of Medical Laboratory Sciences, Faculty of Health Sciences, Bamenda University of Science Technology (BUST), Cameroon; Department of Public Health, Faculty of Health Sciences, Texila American University, Georgetown, Cooperative Republic of Guyana; Department of Biochemistry and Molecular Biology, Faculty of Science, University of Buea, P. O. Box 63 Buea, Cameroon; Infectious Disease Laboratory, Faculty of Health Sciences, University of Buea, P. O. Box 63 Buea, Cameroon; Central African Network for Tuberculosis, HIV/AIDS and Malaria (CANTAM), University of Buea, Buea, Cameroon; Department of Biological Sciences, Faculty of Sciences, University of Bamenda, Cameroon; Dapartment of Medical Laboratory Sciences, Faculty of Health Sciences, University of Bamenda, Cameroon; Departmen of Public Health, Faculty of Health Sciences, University of Information Technology (ICT-U), Yaoundé, Cameroon; Laboratory Department, Holy Family Hospital Akum, Bamenda, Cameroon; Centre for the Study and Control of Communicable Diseases, Faculty of Medicine and Biomedical Sciences, University of Yaoundé I, Yaoundé, Republic of Cameroon; Department of Microbiology, Parasitology, Haematology and Infectious Diseases, Faculty of Medicine and Biomedical Sciences, University of Yaoundé 1, Yaoundé, Republic of Cameroon; IMPM, Ministry of Scientific Research Yaoundé, Cameroon

**Keywords:** Hepatotoxicity, Combined Antiretroviral Therapy (cART), HIV/AIDS, Transaminases, Bali, Cameroon

## Abstract

**Introduction:** The incidence of hepatotoxicity is life-threatening and can result to an end-stage liver disease in long-term patients on combined antiretroviral therapy (cART). Our study sought to evaluate the incidence and predictors of cART-induced hepatotoxicity (CIH) among long term users on cART in a rural District hospital.

**Methods:** This was a hospital-based cross-sectional study in the Bali District Hospital. Spectrophotometric method was use for the quantitative measurement of alanine-aminotransferase (ALT) and aspartate-aminotransferase (AST) levels. Patients with elevations of both ALT and AST were considered CIH. The Chi (χ^2^) square test, ANOVA and Kaplan Meier log-ranked/ survival analyses were used to analyse the data.

**Results:** Of the 350 participants enrolled [156 (44.6%) males and 194 (55.4%) females], aged 43.87 ± 0.79 years (range 20 – 84 years) included in this analysis, 26 (4.4%) experienced moderate CIH. We observed 57 (16.3%), 62 (17.7%) and 238 (68%) elevated levels ALT + AST, ALT and AST respectively. Two independent predictive factors of CIH were, the male sex and alcoholism during the study period.

**Conclusion:** The prevalence of CIH in HIV-infected patients in Bali was lower than that observed in previous studies. The duration of therapy had no influence on the frequency of CIH. Alcoholism and smoking showed significant differences in the development of CIH.

## 1. Introduction

The human immune deficiency virus/ acquired immunodeficiency syndrome (HIV/AIDS) is a devastating infection that remains a public health problem in Sub-Saharan Africa, is caused by either of two lentiviruses: HIV-1 or HIV-2[1, 2].Of the 36.7 million persons living with HIV (PLWHIV), with an estimated 2.1 million new infections, representing a rate of 0.3 new infections per 1,000 uninfected people worldwide, 19 million (51.8%) are in Sub-Saharan Africa[3, 4].In Cameroon, the incidence of adult HIV has fallen consistently from 7.7% in 1999 to 4.3% in 2013 but has remained high among female sex workers, with an estimated prevalence of 3.6% (range; 1.5 – 6.3%) within the ten Regions[5]. Highly Active Anti-Retroviral Therapy (HAART) is a combination of antiretroviral drugs for the management of HIV/AIDS [6, 7].The first-line HAART recommended for use in Cameroon by the Ministry of Public Health (MOH) and the World Health Organization (WHO) areTenofovir /Lamivudine/ Efavirenz(TDF+3TC+EFV)[8, 9].

Although HAART and combination antiretroviral therapy (cART) constitute the most significant interventions that have changed the landscape of HIV-related morbidity and mortality, there are also challenges of adverse drug reactions leading to dose modifications, changes or treatment discontinuations [10–12].

Highly Active Anti-Retroviral Therapy (HAART)-associated hepatoxicity/ cART-induced hepatotoxicity (CIH), arbitrarily defined as AST or ALT > 3 X upper limit of normal (ULN) in the presence of symptoms, or serum AST or ALT > 5 X ULN in the absence of symptoms, usually resolves without modification of therapy [13–18]. Hepatotoxicity, which is the most cited reason for the withdrawal of approved drugs, is damage caused by exposure to a drug or non-pharmacological agents[19],and is consequently associated with HAART/ cART[15, 20] especially nevirapine-based HAART [21] or efavirenz-based drug-induced liver injury [22]. In isolated instances, serious and life-threatening conditions may arise.

Theclinical presentation of HAART/ cART-induced hepatotoxicity can range from mild asymptomatic increases in serum transaminases to overt liver failure [15, 23, 24]. Retrospective studies indicate that, the incidence of cART-related severe hepatotoxicity is approximately 10%, and life-threatening events occur at a rate of 2.6 per 100 person years [25]. The incidence of drug-induced hepatotoxicity in the general African population is estimated to fall between 1/100,000 and 20/100,000 [26]. Hepatotoxicity due to ART may be related to agents from a number of classes including nucleoside reverse transcriptase inhibitors (NRTIs), non-nucleoside reverse transcriptase inhibitors (NNRTIs) and protease inhibitors [27]. The severity of hepatotoxicity may range from transient elevations in transaminase levels to hepatic failure and death, via a variety of mechanisms such as direct cell stress and disturbances in lipid/ sugar metabolism and steatosis, as seen with PI [27]. Co-infection with hepatitis B virus (HBV) or hepatitis C virus (HCV) has consistently been associated with increased risk of ART-related hepatotoxicity [25]. Other risk factors associated with ART-related liver injury include pre-existing advanced fibrosis, pre-treatment of elevated ALT or AST, alcohol abuse, old age, female gender, first exposure to ART, significant increase in CD4^+^ cell count after ART initiation, concomitant tuberculosis medications and cocaine use [28].

While all antiretroviral drugs have some risks of hepatotoxicity, some are more implicated than others. The non-nucleoside reverse transcriptase inhibitors (NNRTI) typically cause either hypersensitivity reactions or direct drug toxicity and therefore have two peaks of onset: within days to weeks or several months after initiation [25].Nevirapine (NVP) is the NNRTI most associated with hepatotoxicity[21], though hypersensitivity reactions resulting in liver failure have been reported with the newer NNRTI etravirine.Efavirenz and stavudine can also cause hepatotoxicity but does so less frequently than NVP or etravirine[22].

Many studies have been carried out in Cameroon [9, 29, 30] and out of Cameroon [14, 15, 23, 31–33] on hepatotoxicity but there is paucity data on HAART/ cART-inducedhepatotoxicity.

The main objective of this study was to determine the incidence of hepatotoxicity and to assess the possible risk factors for developing cART-induced hepatotoxicity among PLWHIV at the Bali District Hospital.

## 2. Mategrial and Methods

A retrospective study of HIV – infected patients was conducted to examine the incidence and predictors of cART-induced hepatotoxicity (CIH).

### 2.1 Study Area

The study was conducted at the state-owned Bali District Hospital in the North West Region of Cameroon between December 2017 to January 2018. Bali District Hospital is found in Bali Sub Division of Mezam Division in the North West Region of Cameroon.

Bali has a population of about 37,103 inhabitants[34],which is made up of mostly indigenouspeople, with a considerable proportion of non-indigenes. The most predominant activity of the people of Bali is farming.

### 2.2 Study Design and Population

A retrospective cross-sectional study was designed to examine the incidence and predictors of CIH. Study participants were Persons living with HIV, who were on cART/ HAART.

Participants aged ≥ 18 years and willingness to have HIV status confirmed from clinical records or by a point-of-care test were included in the study. Pregnant women, those younger than18 years, those on unprescribed medications and severely sick persons were excluded from the study.Participants with CD4^+^> 500 cells/μL and viral load < 500 copies/mL were excluded from this analysis.

The sample size was calculated using the CDC-Epi Info^™^ 7.2.3.1 StatCalc software with the following characteristics: an estimated population size for Bali Health area of 37,103 inhabitants [34], expected frequency of persons living with HIV on ART in Bali of 50%, accepted error margin of 5%, design effect of 1.0 and one cluster. Thus, the CDC-Epi Info^™^ 7.2.3.1 StatCalc estimated minimum sample size was 380.A final sample of 389 participants were enrolled into the study and 350 (89.9%)were included in this analysis.

### 2.3 Data collection tool and Data Collection

The instruments used for the collection of data were, a well-organised laboratory form and patients’ files. Data collected for analysis was defined as; socio-demographic information (age, gender and marital/educational status) and transaminase (ALT and AST) concentrations. The transaminase concentrations were obtained by spectrophotometric measurements.

### 2.4 Specimen Collection and analysis

3 mL of blood specimen was collected from each participant by venipuncture into 5 mL vacutainer dry test tubes and allowed to clot at room temperature. The specimens were latter centrifuged for five minutes at 2500 rpm, and the sera collected were assayed for ALT and AST according to manufacturers’ instructions (Quimica, Italy).

#### Laboratory analysis

All the ALT and AST measurements wereperformed at the Bali District Hospital Laboratory.

Hepatotoxicity was defined based on biochemical measurements as an elevation in serum ALT and/ or AST from the normal following the International Consensus Criteria and also on previous studies [15, 19, 23].The severity of the liver injury was indicated by category (graded) based on various enzyme levels (Grade 1/ Mild, Grade 2/ Moderate, Grade 3/ Severe, and Grade 4/ Acute Liver Failure) [6, 16].

### 2.5 Statistical Analysis

All data collected was entered into epi info 7.2.3.1 for analyses.The variables that presented associationswith the outcome after bivariate analyses were entered intomultivariate logistic regression model to identify independentpredictors of cART-induced hepatotoxicity (CIH). The probabilities of developing hepatotoxicity with duration of antiretroviral treatment were estimated by Kaplan Meier methods and log-ranktest was used to determine statistically significant association.The Statistical Package for the Social Sciences (SPSS), version 25.0 for windows (IBMCorp.released 2017) was used for the multivariate regression analysis as well as the Kaplan Meier survival analysis.

Univariate and multivariate Cox proportional hazards regressions, were performed to assess the predictors of cART-induced hepatotoxicity. The variables included in the multivariate model were those with either a theoretical importance or ones with a *p*<0.05 in theunivariable models.

### 2.5 Ethical considerations

The study was approved by the Regional Delegate of Public Health for the North West Region and the Higher Institute of Health Sciences of Bamenda University of Science and Technology (BUST) and was conducted in accordance with the Helsinki declaration [35]. An administrative clearance was obtained from the Director of the Bali District Hospital.All participants signed informed consent forms and all records were strictly confidential.

## 3.Results

### Socio-Demographic Characteristics

A total of 350 patients [156 (44.6%) males and 194 (55.4) females] aged 43.87 ± 0.79 years (range 20 – 84 years), with a mean (± SEM) duration of cART of 6.30 ± 0.21 years (range 0 – 13 years), were included for analysis in this study. A majority 154 (44%) of the study participants were in the age group 40 - < 60 years old and most of them 148 (42.3%) had at least the secondary education (Table 1).

**Table 1:**
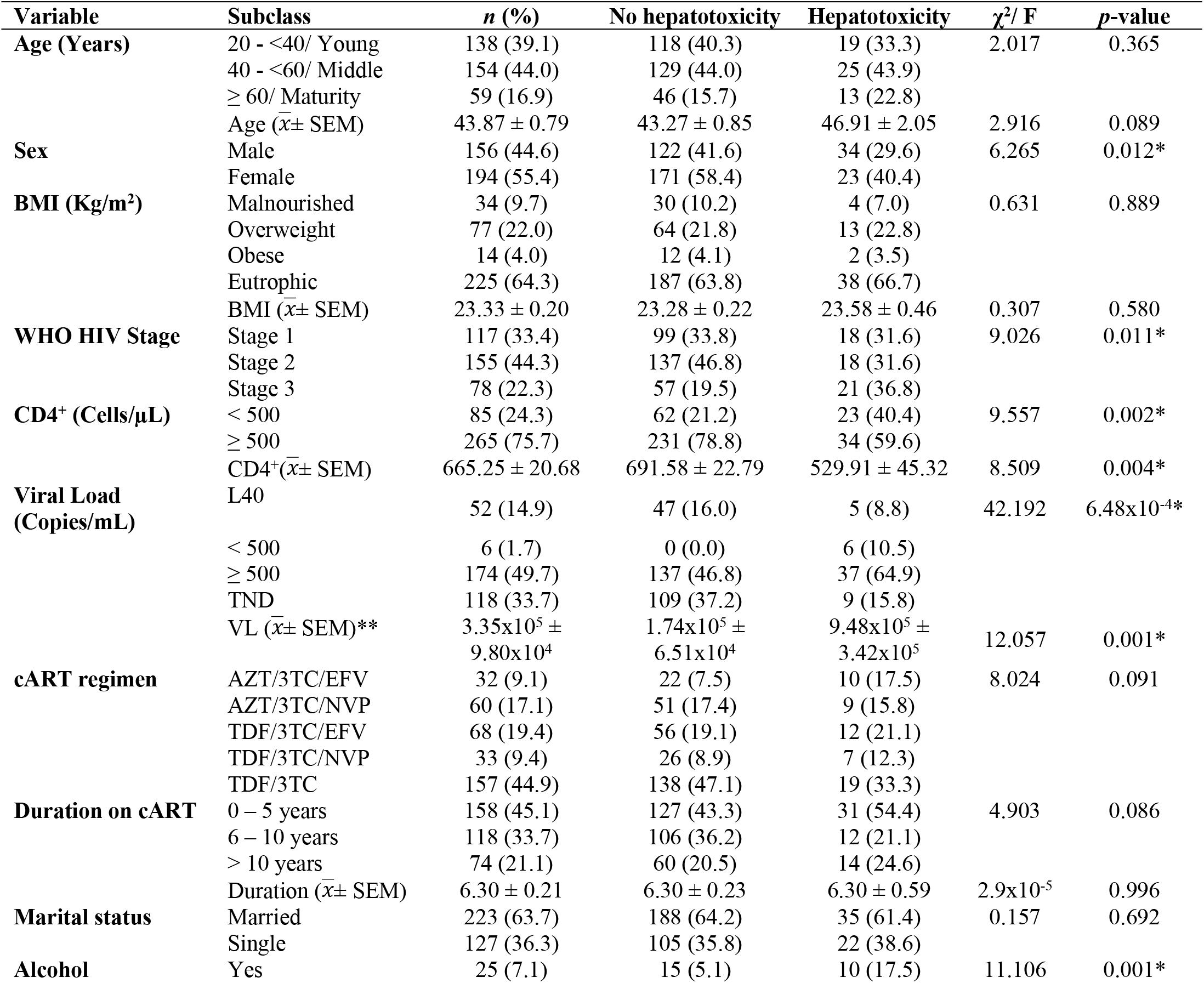
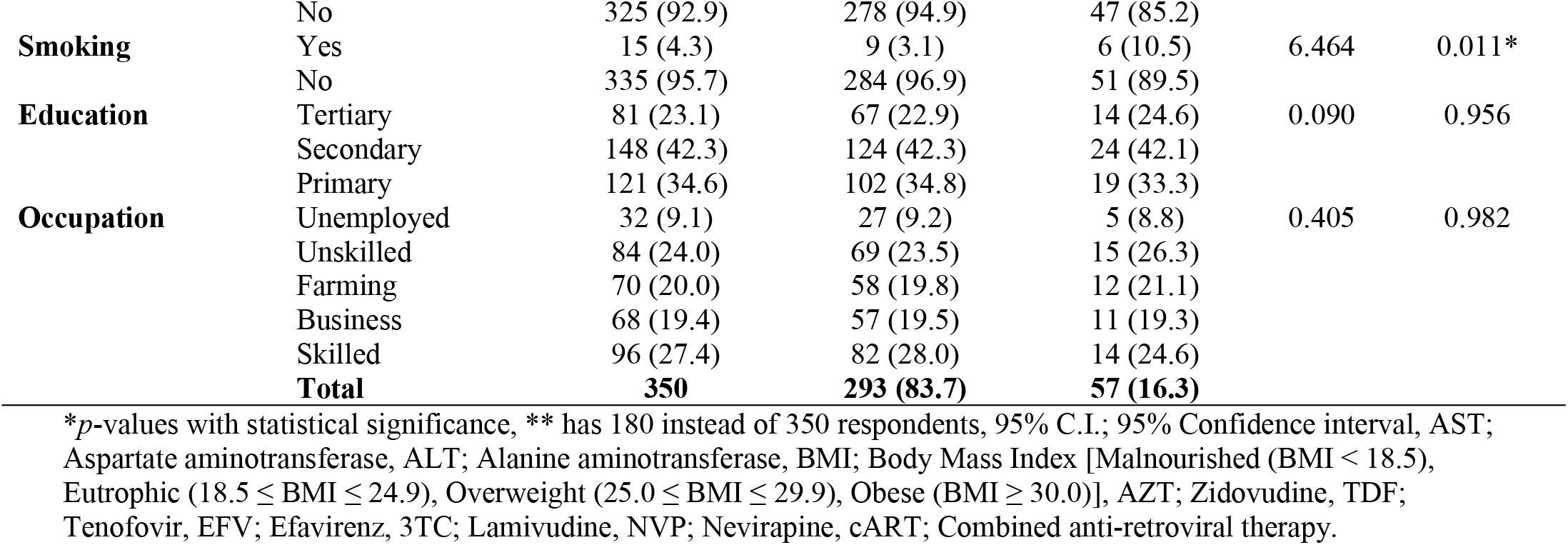
Socio-demographic and Clinical Characteristics of patients with and without hepatotoxicity.

Thirty-four males (29.6%, N = 57) and 23 females (40.4%, N = 57) among cases had cART-induced hepatotoxicity. Of the 350 study participants, 34 (9.7%%) were malnourished as defined by body mass index (BMI) of < 18.5 Kg/m^2^, 77 (22.0%) were overweight, 14 (4.0%) were obese and 64.3% were eutrophic (normal). Most of the participants 223 (63.7%) were married and just one-fifth (20.0%) of them were farmers (Table 1).

There was no significant difference as regards age, BMI, cART regimen and duration of therapy between cases and controls.

### Clinical and Biochemical Spectrum of Participants

Majority 155 (44.3%) of the study participants had WHO stage 2 HIV, 117 (33.4%) had stage 1, while theremaining 78 (22.3%) had stage 3 HIV.The mean (± SEM) CD4^+^ count was 529.91 (± 45.32) and 691.58 (± 22.79) for cases and controls, respectively.

Ten alcoholics (17.5%, N = 57) and six smokers (10.5%, N = 57) were among the cases who had CIH.

The mean (± SEM) values of ALT and AST were 30.18 (± 0.76)U/L and 49.52 (± 1.06)U/L and were significantly higher amongst cases [55.00 (± 1.82)U/L vs 72.61 (± 3.43)U/L] than controls [25.35 ± (0.46)U/L vs 45.03 (± 0.85)U/L] (Table 2).

**Table 2:**
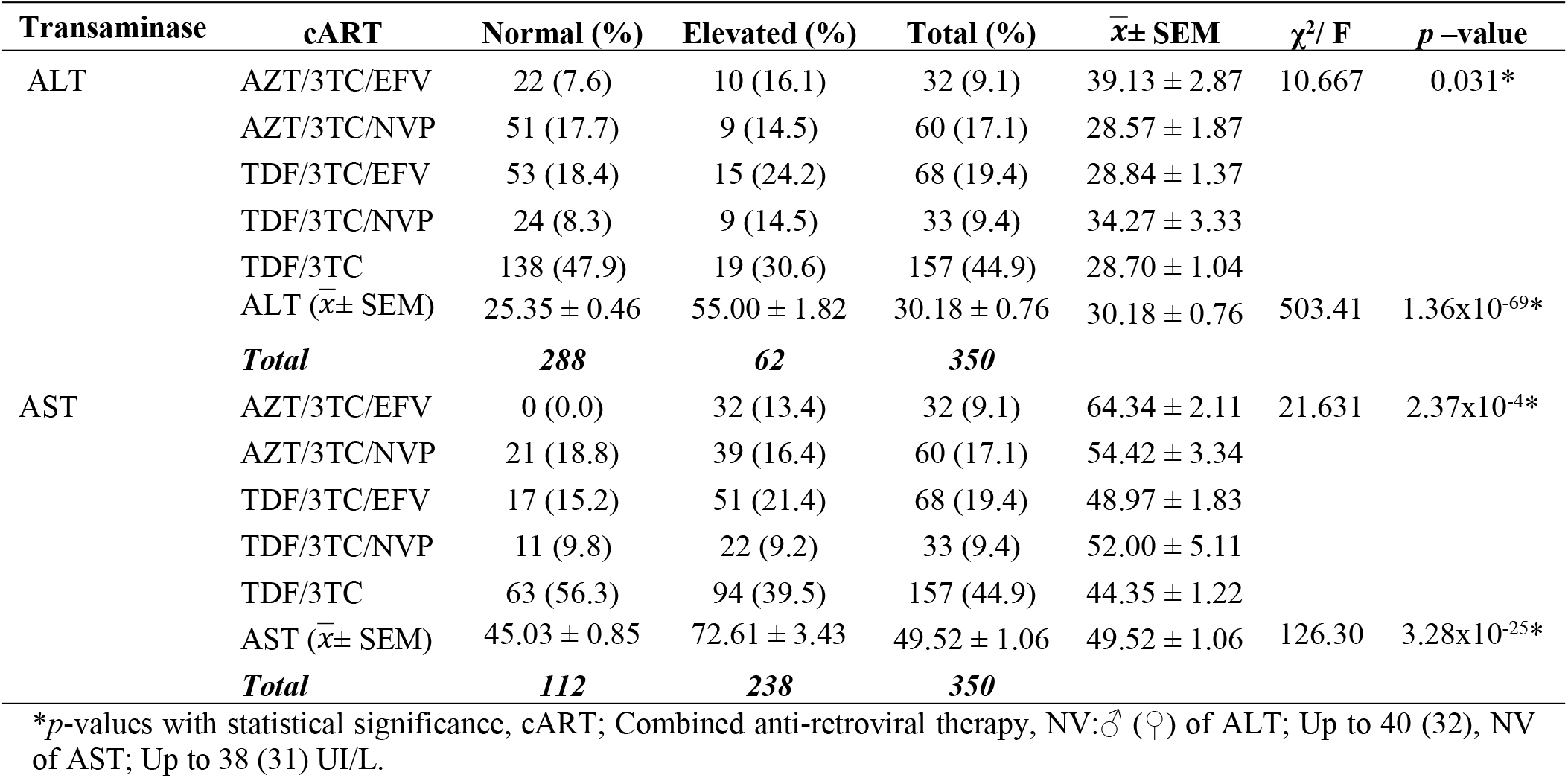
Bivariate analysis of cART association withelevated transaminases.

Table 2 shows the prevalence of elevated liver enzymes in the five cART treatment groups. Patients on TDF/3TC, had the highest incidence19 (30.6%) and 94 (39.5%) of elevated ALT and AST respectively, followed by TDF/3TC/EFV with incidence of 15 (24.2%) and 51 (21.4%) of elevated ALT and AST respectively. However, the overall prevalence of hepatotoxicity in the five treatment groups for the two liver enzymes was 57 (16.3%).

In all patients with cART-induced hepatotoxicity, 31 (8.9%) had mild hepatoxicity while 26 (7.4%) had moderate hepatotoxicity. Alanine aminotransferase (ALT) related hepatotoxicity 62/350 (17.7%) was lower compared to AST related hepatotoxicity 238/350 (68%).Alanine aminotransferase and AST hepatotoxicity were significantly (*p* = 0.031 vs *p* = 2.37×10^−4^) associated with cART regimens (Table 2).

Results of the study indicated that there were five different cARTregimens (Figure 1; Table 1). A majority 157 (44.9%) of the study participants were on TDF/3TC, followed by 19.4% of TDF/3TC/EFV, 17.1% of AZT/3TC/NVP, 9.4% of TDF/3TC/NVP and the least was 9.1% of AZT/3TC/EFV.

**Figure 1:**
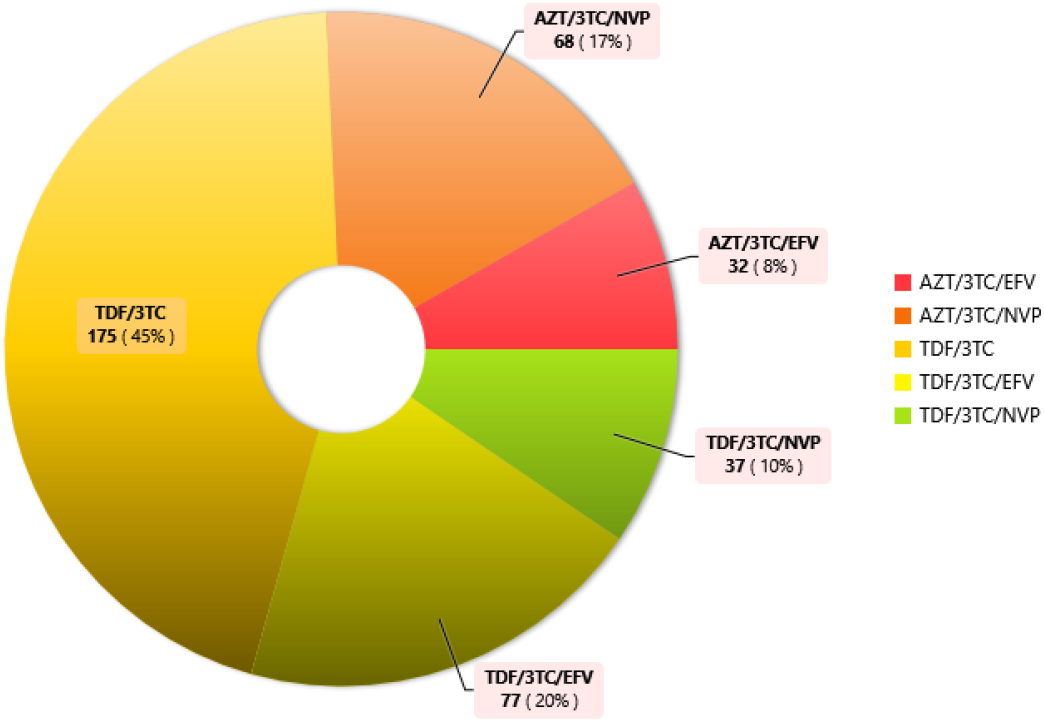
Regimens of combined ART.

### Factors associated with cART-induced hepatotoxicity

Age was categorized into three groups and there was no difference betweencases and controls between these age groups. The nutritional status (as assessed by BMI) of study participants seems to be good [mean (± SEM), BMI; 23.33 (± 0.20) kg/m^2^].Cases were less likely to have malnutrition as compared to controls, 4 (7.0%) vs 30 (10.2%), respectively; (*p*= 0.889) (Table 1). The mean BMI was almost the same among cases as compared to controls, 23.58 (± 0.46) kg/m^2^vs 23.28 ± 0.22 kg/m^2^, respectively. The proportion of patients with WHO clinical stage 2 HIV was higher in the controlgroup137 (46.8%, N = 293)while in the cases group the number of WHO clinical stage 2 HIV was 18 (31.6%, N = 57) (*p* = 0.011).Cases were also more likely to have lower CD4^+^ count as compared to controls; 23 (40.4%, N = 57) of cases and 62 (21.2%, N = 293) of controls had CD4^+^ count less than 500 cell/ μL (*p* = 0.002). Therefore;sex, WHO clinical stagesof HIV/AIDS, CD4^+^/VL, alcoholism and smoking were significantly associated withcART-induced hepatotoxicity from bivariate model analysis (Table 1).

Univariate analysis was also done using Cox Proportional hazard regression analysis for the variables in Table 3. Sex and alcohol consumption were observed to be predictors of cART-induced hepatotoxicity. In the multivariate model, the predictors of developing cART-induced hepatotoxicity include the male sex (Hazards Ratio (HR) = 1.6, 95% C.I = 0.9 – 2.8), the cART regimen AZT/3TC/EFV [HR = 2.5, 95% C.I = 1.1 – 5.7] and alcohol consumption [HR = 15.3, 95% C.I = 6.3 – 37.1].

**Table 3:**
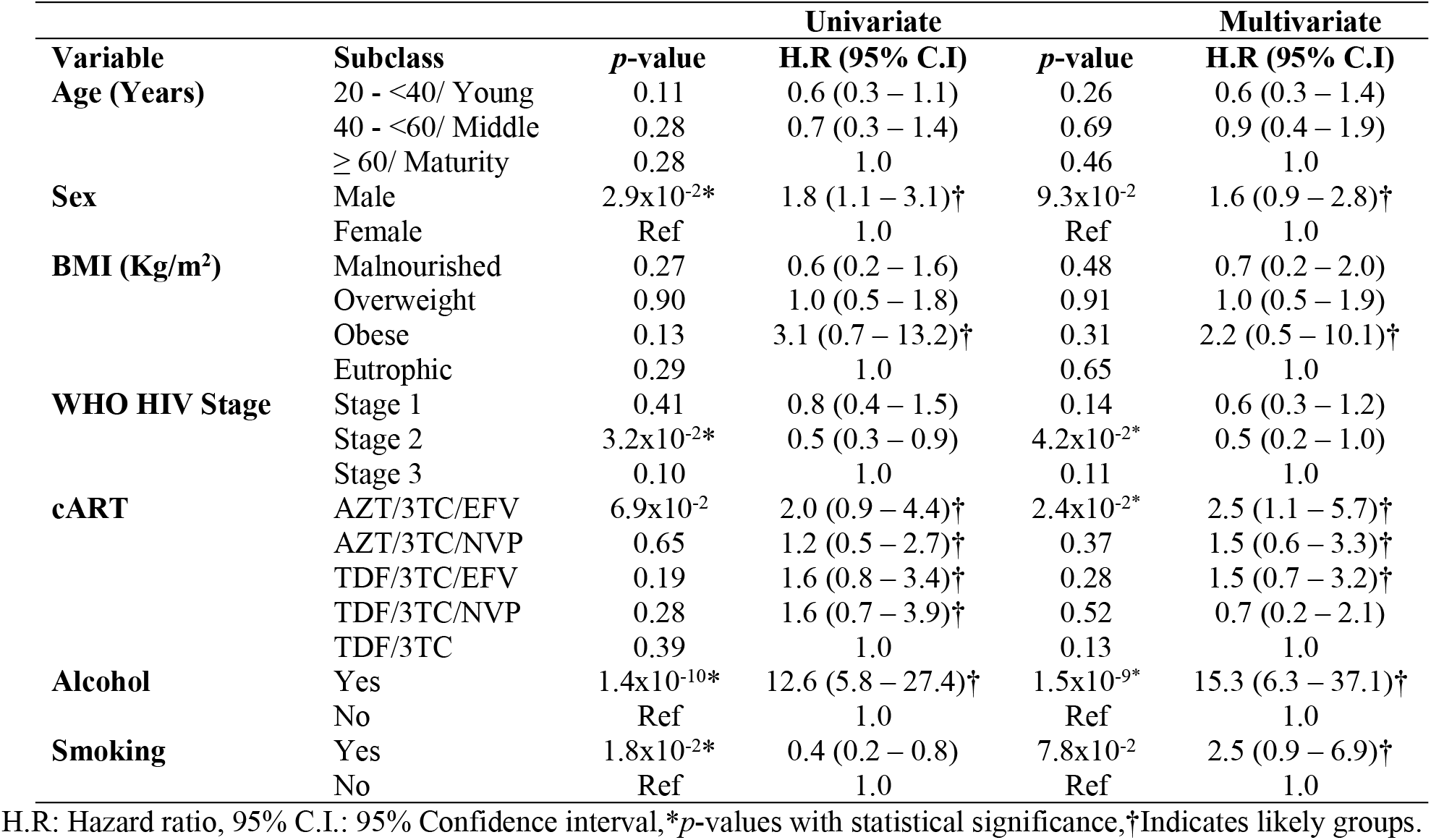
Univariate and Multivariate Cox proportional regression analysis to show the risk factors for developingcART-induced hepatotoxicity.

The mean (±SEM) time to onset of cART-induced hepatotoxicity was lower among patients with WHO clinical stage 3 HIV as compared to WHO clinical stage 2 and WHO clinical stage 1; 2.61 (±0.09), 2.83 (±0.05) and 2.78 (0.05) patients, respectively (χ^2^ = 7.108; *p* = 0.029). Kaplan Meier analysis shows that patients with obesity (BMI > 30 kg/m^2^) were likely to develop hepatotoxicity within 1.86 (±0.94) mean years while the malnourished, overweight and eutrophic patients were likely to develop hepatotoxicity in 2.89 (±0.08), 2.73 (±0.08) and 2.76 (±0.04) mean years of initiation of cART (χ^2^ = 1.4818; *p* = 0.701) (Figure 2).

**Figure 2:**
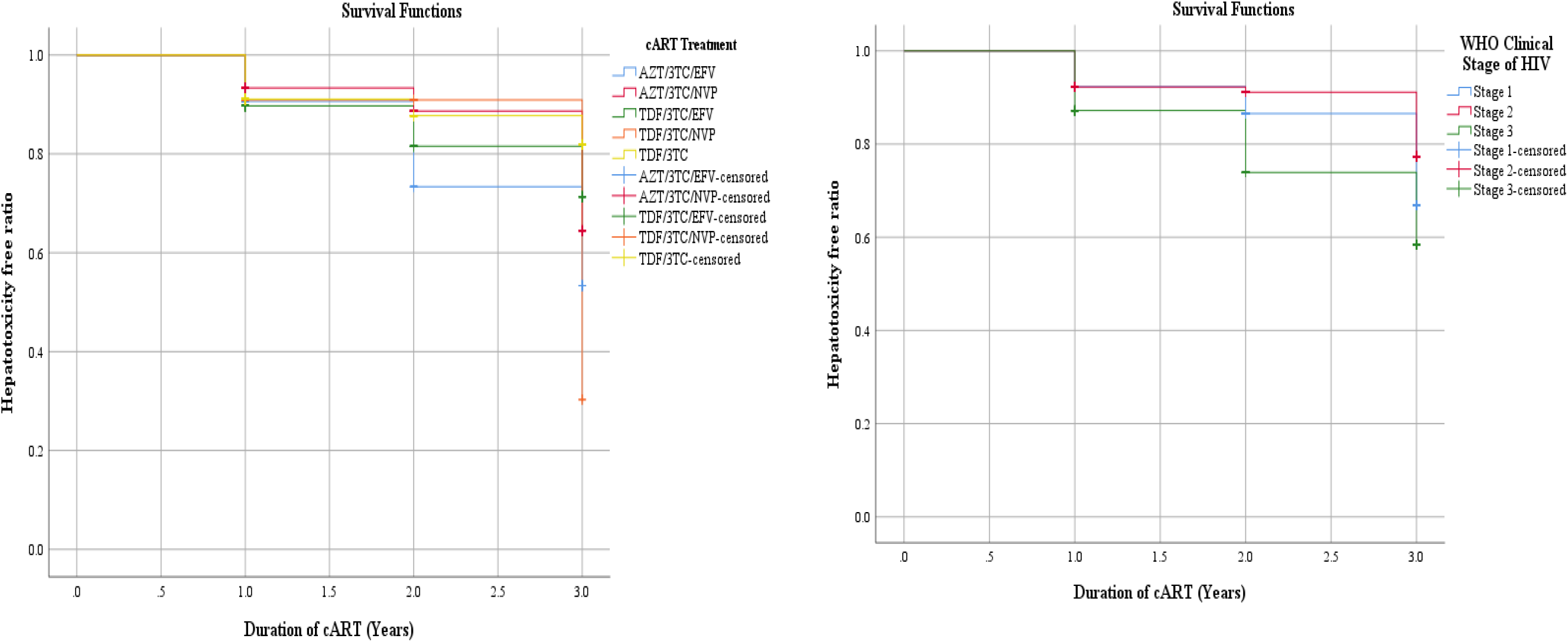
Kaplan-Meier Survival curves of hepatotoxicity with respect to cART treatment groups and WHO Clinical stage.

The mean (±SEM) time to onset of cART-induced hepatotoxicity among patients on the different treatments were 2.64 (±0.13), 2.82 (±0.07), 2.71 (±0.08), 2.82 (±0.11) and 2.78 (±0.05) respectively for AZT/3TC/EFV, AZT/3TC/NVP, TDF/3TC/EFV, TDF/3TC/NVP and TDF/3TC. Comparing the survival curves of hepatotoxicity, we observed that there is no statistically significant difference in occurrence rates of the participants in the various treatment groups and that the grouping has no significant influence on time of onset (χ^2^ = 5.054; *p* = 0.282) (Figure 2). The overall mean Survival Proportion (SP) and Standard Error (SE) with respect to the years of treatment were: SP = 2.76 and SE = 0.034.

## 4. Discussion

### General Characteristics

Hepatotoxicity is one of the most common adverse drug reactions associated with HAART/cART in Persons Living withHIV/AIDS. This increases the mortality rate in PLWHIV, rather than the HIV infection itself.

As observed in this study, age difference is not a determinant factor for ALT or AST elevations.This is supported by studies reported in Africa [14, 36] and Zürich-Switzerland [37]which reported that age is not a risk factor for the development of hepatotoxicity in patients on cART/HAART.However, studies carried out elsewhere in Africa reported that age was significantly associated with drug-induced hepatotoxicity[15, 31]. These differences could be due to the fact that more than 60% of our study participants were above 40 years of age.

In our study, many more femalesdevelopedhepatotoxicitywhen compared with males (Table 1).Our findings are similar to those of studiesreported in Fako-Cameroon[29, 30], elsewhere in Ethiopia[15, 31, 33] and in Milan-Italy[21].However, studies in Zürich-Switzerland [37], Tanzania [36] and Ethiopia [23], reported higher rates of hepatotoxicity amongmale than female patients on cART. These differences could be as a result of the fact that this study, weenrolled only HIV patients.

### Hepatotoxicity

In the present study, the overall cART-induced hepatotoxicity among HIV-infected patients was 16.3%, which waslower compared to the 42.4 – 54% reported in Cameroon[29, 38],20.1 – 32% in Ethiopia [23, 32, 39], 25% reported in Warsaw-Poland [12]and 19.7/100 person years in South Africa[28]. Our findings were higher compared to the 13.6% reported in Fako-Cameroon[30], 11.5% in Ethiopia [33] and 7.8% in Tanzania[36]. However, our study is consistent with the 16% reported in Zürich-Switzerland[37]and Taiwan-China [40]and the 15 – 16.7%reported in Ethiopia[15, 31]. This could be explained by the fact that the present study did not include tuberculosis and tuberculosis/ HIV co-infected patients, neither did it include hepatitis B and C viral infections as well as co-infections.

The finding of this study indicated that ALT and AST related hepatotoxicity were 17.7% and 68% respectively. This was in line with the findings of a study reported in Ghana[14] and lower when compared to findings reported in Yaoundé-Cameroon [38].

### Predictors of hepatotoxicity

We observed a relatively higher incidence of hepatotoxicity amongst case patients receiving TDF/3TC/EFV (21.1%, N = 57), AZT/3TC/EFV (17.5%, N = 57) and TDF/3TC/NVP (12.3%, N = 57) compared to those on other treatment groups. Although there was no statistical association of cART treatment groups with hepatotoxicity, there were significant variations of the elevated transaminases with the cART treatment groups (Table 2). The highest mean 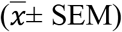 elevations of both ALT (39.13 ± 2.87) and AST (64.34 ± 2.11) were observed in TDF/3TC/EFV treatment group (Table 2). These elevations were similar to those reported in Cameroon [29, 38] and in Ghana [14].

In this study, it was observed that the sex (male) and alcohol consumption were the independent predictors of cART-induced hepatotoxicity. The WHO clinical stage 2 of HIV, AZT/3TC/EFV regimen and cigarette smoking were significantly associated with hepatotoxicity (Table 3). Our finding of male being a predictor was in line with that of a study reported in Ethiopia [32] and different from females as reported in Ethiopia [23, 31]. Alcoholism as reported in this study was in line with alcoholism reported in London [16], Zürich [37], as well as concomitant administration of other drugs [15, 17, 28].

## 5. Strengths and limitations of the study

Strengths of the study: The data used was collected by experienced scientists, using laboratory forms and patients’ records.

Limitations of the study: A limitation to the study was that it was a cross-sectional study collecting data on the dependent and independent variables at the same time. It equally did not consider the concomitant administration of other therapeutic agents; anti-malaria drugs, herbs, as well as smoking and alcohol intake.

## 6. Conclusion

In conclusion, cART-induced hepatotoxicity is incident amongst HIV-infected patients seeking health care in the Bali District Hospital, regardless of the cART treatment group. Allof these hepatotoxic events were not severe and had no clinical significance. The male sex and alcoholism were associated with a higher rate of liver injury. Prospective studies focusing on the effects of cART/ HAART on hepatitis in HIV-infected patients are needed to confirm our findings.Combined antiretroviral therapy is frequent and a major concern amongst Cameroonian HIV patients and regular monitoring of liver enzymes during early therapy is recommended for proper identification and management of cART-induced hepatotoxicity.

## Declarations

### Ethical approval and consent to participate

Ethical clearance was obtained from the North West Regional Delegation of Health and the Higher institute of Health Sciences of Bamenda University of Science and Technology (BUST)and was conducted in accordance with the Helsinki declaration [35]. An administrative clearance was obtained from the Director of the Bali District Hospital. Participation in the study was voluntary, and all participants signed informed consent.

## Abbreviations

3TC: Lamivudine, 95%
C.I: 95% Confidence Interval
AIDS: Acquired Immune Deficiency Syndrome
ALT: Alanine Amino Transferase
ARV: Antiretroviral Therapy
AST: Aspartate Amino Transferase
AZT: Zidovudine
cART: Combined Antiretroviral Therapy
CIH: cART-Induced Hepatotoxicity
CD4^+^: Cluster of Differentiation-4^+^ cells
EFV: Efavirenz
HR: Hazard Ratio
HA: Health Area
HAART: Highly Active Antiretroviral Therapy
HIV: Human Immunodeficiency Virus
N(NRTI): Non (Nucleoside reverse transcriptase inhibitor)
OR: Odds Ratio
*p*: Significance value
VL: Viral Load
WHO: World Health Organisation
*χ*^*2*^: Chi square.

## Acknowledgments

Special thanks to our study participants, without whom this study would have been fruitless. We cannot underestimate the collaboration of our friends, colleagues and the Laboratory staff of Bali District Hospital, Bamenda, Cameroon for their immense support towards the realization of this work.

## Authors’ contributions

YEN, FNC and NI conceived and designed the study; YEN, BSE, NGF and FN enrolled patients, collected specimens and performed the experiments; YEN and FNC curated the data and performed the statistical analyses; YEN, FNC and NI searched for literature and wrote the first draft of the manuscript; FNC, GMI and NI supervised the study; YEN and NI provided reagents, materials and analysis tools; YEN, FNC, GMI contributed to the discussion and scientific content; All authors contributed to the write up, reviewed the final draft, read and approved the final manuscript.

